## Supplementary Information for "Modelling genetic stability in engineered cell populations"

### Modelling genetic stability in engineered cell populations: supplementary information

### S1 Supplementary model implementation

#### S1.1 Host-aware model components

The general model for our framework comprises one set of Ordinary Differential Equations (ODEs) for each cell phenotype considered, as is also given in Equation (5) of the main text:

$$\text{Mutation-aware model} \triangleq \{\dot{X}_1, \dot{\Psi}_{\mathrm{X}_1}, \ldots, \dot{X}_n, \dot{\Psi}_{\mathrm{X}_n}\}. \tag{S1}$$

Each set includes one ODE that describes the number of cells in the considered state ($\dot{X}_i$), and a set of ODEs ($\dot{\Psi}_{\mathrm{X}_i}$), based on the model of Weisse et al. [1], that describes the host-aware growth dynamics of the cells in that state. The equations in $\dot{\Psi}_{\mathrm{X}_i}$ describe the dynamics of 13 variables when one heterologous protein species is modelled: energy ($e$), proteins of four different classes (ribosomal ($R$), enzymatic ($Z$), housekeeping ($Q$) and heterologous ($H$)), mRNAs for each class ($m_R$, $m_Z$, $m_Q$, $m_H$), and mRNA-ribosome complexes for each class ($c_R$, $c_Z$, $c_Q$, $c_H$). In Weisse et al.'s model, the proteins for energy metabolism are further split into those for transporting nutrients from outside to inside the cell and those for metabolising intracellular nutrients into energy. As we are not concerned about modelling the effects of nutrient limitation, we simplify these steps by combining 'transport' and 'metabolism' into one enzymatic ($Z$) class and adjust reaction rates accordingly (Table S1). In full, the set of host-aware ODEs for a general cell state can be written as:

$$\dot{\Psi} = \begin{cases} \dot{e} = \epsilon(Z) - \sum_x (\Gamma_x(c_x, e)) - \lambda \cdot e, & \text{(S2)} \\ \dot{R} = \Gamma_R(c_R, e) + \sum_x (k_x^- \cdot c_x) - \sum_x \left(k_x^+ \cdot m_x \cdot R + \Gamma_x(c_x, e)\right) - \lambda \cdot R, & \text{(S3)} \\ \dot{Z} = \Gamma_Z(c_Z, e) - \lambda \cdot Z, & \text{(S4)} \\ \dot{H} = \Gamma_H(c_H, e) - \lambda \cdot H, & \text{(S5)} \\ \dot{Q} = \Gamma_Q(c_Q, e) - \lambda \cdot Q, & \text{(S6)} \\ \dot{m}_R = \omega_R(e) - k_R^+ \cdot m_R \cdot R + k_R^- \cdot c_R + \Gamma_R(c_R, e) - (\lambda + d_m) \cdot m_R, & \text{(S7)} \\ \dot{m}_Z = \omega_Z(e) - k_Z^+ \cdot m_Z \cdot R + k_Z^- \cdot c_Z + \Gamma_Z(c_Z, e) - (\lambda + d_m) \cdot m_Z, & \text{(S8)} \\ \dot{m}_Q = \omega_Q(e) - k_Q^+ \cdot m_Q \cdot R + k_Q^- \cdot c_Q + \Gamma_Q(c_Q, e) - \lambda \cdot m_Q, & \text{(S9)} \\ \dot{m}_H = \omega_H(e) - \beta \cdot m_H \cdot R + k_H^- \cdot c_H + \Gamma_H(c_H, e) - (\lambda + d_m) \cdot m_H, & \text{(S10)} \\ \dot{c}_R = k_R^+ \cdot m_R \cdot R - k_R^- \cdot c_R - \Gamma_R(c_R, e) - \lambda \cdot c_R, & \text{(S11)} \\ \dot{c}_Z = k_Z^+ \cdot m_Z \cdot R - k_Z^- \cdot c_Z - \Gamma_Z(c_Z, e) - \lambda \cdot c_Z, & \text{(S12)} \\ \dot{c}_Q = k_Q^+ \cdot m_Q \cdot R - k_Q^- \cdot c_Q - \Gamma_Q(c_Q, e) - \lambda \cdot c_Q, & \text{(S13)} \\ \dot{c}_H = \beta \cdot m_H \cdot R - k_H^- \cdot c_H - \Gamma_H(c_H, e) - \lambda \cdot c_H. & \text{(S14)} \end{cases}$$

When multi-gene synthetic constructs are being modelled, three additional ODEs are required for each gene. The additional ODEs capture (i) the protein ($H$), mRNA ($m_H$), and mRNA-ribosome complex ($c_H$) for each gene of the synthetic construct. In these cases, variables belonging to the same class are differentiated with subscripts, for example $H_1$ and $H_2$. In the equation set $\dot{\Psi}$, the following notations are used:

- $x$ represents a general proteome class such that $x \in \{R, Z, Q, H_1, \cdots H_n\}$, where $n$ denotes the number of heterologous classes considered. Using this notation, we represent $k_H^+$ using the distinct symbol $\beta$ as it is a key parameter for the design of synthetic constructs;
- $\epsilon(Z) = n_q \cdot Z \cdot \frac{\nu_e \cdot s}{K_e + s}$ is the rate of energy metabolism;
- $\Gamma_x(c_x, e) = \frac{c_x \cdot \gamma(e)}{n_x} = \frac{c_x \cdot \frac{\nu_\gamma \cdot e}{K_\gamma + e}}{n_x}$ is the rate of translation elongation for protein class $x$ where $n_x$ is the number of amino acids in protein class $x$;
- $\omega_i(e) = \frac{\nu_{\omega_i} \cdot e}{K_{\omega_i} + e}$ is the transcription rate for protein class $i$ when $i \in \{R, Z, H\}$. Using this notation, we represent $\nu_{\omega_H}$ using the symbol $\alpha$ as it is a key parameter for the design of synthetic constructs;

**Table S1: Parameters used in the mutation-aware model.** 'R', 'Z', 'Q', 'H': proteome classes for 'ribosomal', 'enzymatic', 'housekeeping' and 'heterologous'. 'HSC': half-saturation constant. 'aa': amino acids. 'molecs': molecules.

**a**: can be varied to reflect different designs of synthetic construct. $\alpha_{\mathrm{X}}$ is varied logarithmically to reflect ranges seen in typical promoter libraries; $\beta_{\mathrm{X}}$ is kept constant at same value as the mRNA-ribosome binding rate for the other proteome fractions, but can be varied to simulate different RBS strengths; $n_H$ is kept constant at the average length of cytosolic proteins in *E. coli* [2], but can be varied to simulate different synthetic protein sizes; $z_{\mathrm{X}}$ is varied logarithmically to reflect ranges of mutation rates generated by different sequences (see sources explored in [3]).

**b**: value obtained from using a degradation time of 7 min: $60 \cdot \frac{\ln 2}{7} \approx 5.94$.

**c**: to provide steep auto-inhibition.

**d**: near the diffusion limit.

**e**: $K_e$ and $s$ are chosen relative to each other to create variation in energy metabolism dynamics when $Z$ varies. In [1], $s$ is used instead as a variable with an additional parameter denoting external nutrient quantity.

**f**: parameter fit in [1] using an adaptive Markov Chain Monte Carlo method [4].

**g**: can be varied to reflect different growth conditions. $n_q$ is kept constant at 1, but can be varied to simulate growth media with different concentrations; $N$ is kept constant at $10^9$ to resemble a dilute population of *E. coli* cells where space is non-limiting, but can be varied to simulate turbidostats with different cell density thresholds.

**h**: calculated by combining the maximal rates for energy transport and energy metabolism from [1]'s model as:
$1 \div \left(\frac{1}{43560} + \frac{1}{348000}\right) = 38\,714\,\mathrm{h}^{-1}$.

| Symbol | Name | Value | Unit | Source |
|---|---|---|---|---|
| $\alpha_{\mathrm{X}}$ | Maximum rate of H-transcription in state X | various | molecs $\mathrm{h}^{-1}$ $\mathrm{cell}^{-1}$ | [5][a] |
| $\beta_{\mathrm{X}}$ | mRNA-ribosome binding rate for H in state X | 60 | cell $\mathrm{h}^{-1}$ $\mathrm{molecs}^{-1}$ | **a** |
| $d_m$ | mRNA-degradation rate | 5.94 | $\mathrm{h}^{-1}$ | [6][b] |
| $h_Q$ | Hill coefficient for Q-inhibition | 4 | none | **c** |
| $k^{+}_{R,Z,Q}$ | mRNA-ribosome binding rate for R,Z,Q | 60 | cell $\mathrm{h}^{-1}$ $\mathrm{molecs}^{-1}$ | **d** |
| $k^{-}_{R,Z,Q,H}$ | mRNA-ribosome unbinding rate for R,Z,Q,H | 60 | $\mathrm{h}^{-1}$ | **d** |
| $K_e$ | HSC for energy metabolism | 1000 | molecs | **e** |
| $K_Q$ | HSC for Q-inhibition | 152000 | molecs $\mathrm{cell}^{-1}$ | **f** |
| $K_{\omega_R}$ | HSC for R-transcription | 427 | molecs $\mathrm{cell}^{-1}$ | **f** |
| $K_{\omega_{Z,Q,H}}$ | HSC for non-R-transcription | 4.38 | molecs $\mathrm{cell}^{-1}$ | **f** |
| $K_\gamma$ | HSC for translation | 7 | molecs $\mathrm{cell}^{-1}$ | **f** |
| $M$ | Cell mass | $10^8$ | aa | [2] |
| $n_H$ | Length of H-proteins | 300 | aa $\mathrm{molecs}^{-1}$ | **a** |
| $n_q$ | Nutrient quality | 1 | none | **g** |
| $n_R$ | Length of R-proteins | 7459 | aa $\mathrm{molecs}^{-1}$ | [7] |
| $n_{Z,Q}$ | Length of Z,Q-proteins | 300 | aa $\mathrm{molecs}^{-1}$ | [8] |
| $N$ | Cell capacity of turbidostat | $10^9$ | none | **g** |
| $s$ | Internal nutrient quantity | $10^4$ | molecs | **e** |
| $v_e$ | Maximum rate of energy metabolism | 38700 | $\mathrm{h}^{-1}$ | [9, 10][h] |
| $v_{\omega_Q}$ | Maximum rate of Q-transcription | 56940 | molecs $\mathrm{h}^{-1}$ $\mathrm{cell}^{-1}$ | **f** |
| $v_{\omega_R}$ | Maximum rate of R-transcription | 55800 | molecs $\mathrm{h}^{-1}$ $\mathrm{cell}^{-1}$ | **f** |
| $v_{\omega_Z}$ | Maximum rate of Z-transcription | 248.4 | molecs $\mathrm{h}^{-1}$ $\mathrm{cell}^{-1}$ | **f** |
| $v_\gamma$ | Maximum rate of translation | 72000 | aa $\mathrm{h}^{-1}$ $\mathrm{molecs}^{-1}$ | [2] |
| $z_{\mathrm{X}}$ | Probability of mutating into state X | various | none | **a** |

- $\omega_Q(e) = \frac{\nu_{\omega_Q} \cdot e}{K_{\omega_Q} + e} \cdot \frac{1}{1 + \left(\frac{Q}{K_Q}\right)^{h_Q}}$ is the transcription rate for protein class $Q$ to reflect the fact that $m_Q$ is auto-inhibited.

As in Weisse et al.'s model, all variables are measured in molecules per cell. For bimolecular reaction rates that depend on the concentration of molecular species, we convert units to number of molecules by assuming a fixed cellular volume of 1 µm$^3$, approximately that of *E. coli* in exponential phase [11]. The units and values of the parameter values are given in Table S1, including any differences from the values used in Weisse et al.'s model. The cell's growth rate ($\lambda$) is defined as the total rate of protein production divided by the cell's mass ($M$), which is the combined amino acid weight of all protein species. At steady state, this value is assumed to be a fixed value of $10^8$ amino acids (unit: aa) [2], giving:

$$M = \sum_x (n_x \cdot x) + n_R \cdot \sum_x c_x = 10^8 \,\mathrm{aa}, \tag{S15}$$

$$\lambda = \frac{\gamma(e) \cdot \sum_x c_x}{M}. \tag{S16}$$

#### S1.2 Algorithm for state transitions

The ODE for the number of cells in any state is automatically generated by recording which states are upstream and downstream of that state and implementing the corresponding transition dynamics between them. In turn, this allows mutation frameworks for any number of 'dimensions' and 'states per dimension' to be simulated. In addition, each state requires a different combination of key mutation parameter values (such as maximum transcription rate $\alpha_{\mathrm{X}}$) to reflect its cells' mutation phenotypes. These are automatically generated for each state and are allocated in the correct order by assigning sequential coordinates to each state. These concepts are illustrated in Figure S1 and are detailed below.

(i) The model requires a user to specify the following as an input: the number of mutation states that each mutable part can take ($s$); the number of parts considered for mutation ($d$); a vector of values indicating the probability of each part sustaining a particular mutation ($[z]$); a vector of values indicating the functionality of each part when in different mutation states ($[p]$, part values), such as the maximum transcription rate for E-, I- and M-cells. (ii) Using $s$ and $d$, an algorithm is used to create coordinates for each state in the framework. These are designed such that each state's coordinate has $d$ integers, and each integer's value is between 0 and $s-1$, inclusive. Lower integer values are given to states with higher mutation severities, such that

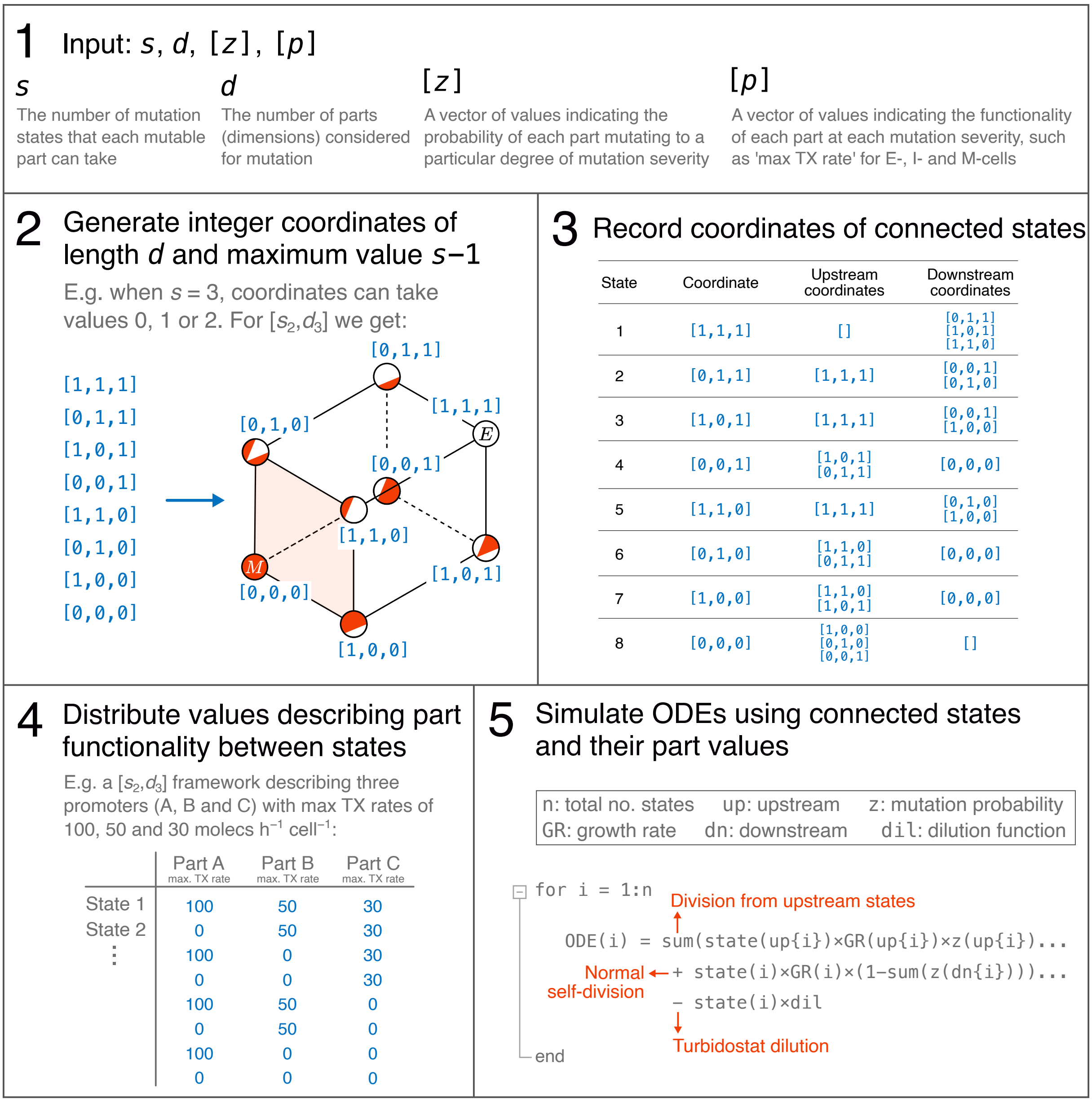


**Figure S1: How the general modelling framework is constructed from a user's input.** The code used to construct any mutation-modelling framework can be split into five steps: (i) user input, (ii) coordinate generation, (iii) recording connections between states, (iv) distributing part values, and (v) simulating ODEs. The details of each step are outlined in the text of Section S1.2. TX: transcription.

the fully-mutated state's coordinate is all zeros. In Figure S1, an example of the coordinate allocation is shown for the $[s_2, d_3]$ framework, where three parts are modelled as either fully active or fully inactive. (iii) For each state, the coordinates in states directly upstream and downstream are determined. (iv) Using the information given in $[p]$, an algorithm is used to distribute the part values to each state in the correct order. For example, with the $[s_2, d_3]$ framework in Figure S1, state 2 (coordinate [0,1,1]) requires its first part to be inactive (0) and its other two parts to be active (1). (v) Finally, the information from (iii) and (iv) are used to create the ODE for the number of cells in each state, which requires values for the growth rates, mutation probabilities ([$z$]) and part values ([$p$]) of the connected states.

#### S1.3 Adjustments to implement the genetic toggle switch

The synthetic toggle switch consists of two mutually repressing genes that express proteins with concentrations $H_1$ and $H_2$. We include inhibitor species $\mathrm{I}_1$ and $\mathrm{I}_2$ with concentrations $I_1$ and $I_2$ that negatively regulate the repression of $H_1$ and $H_2$ respectively. Both synthetic genes require ODEs for their corresponding protein ($H$), mRNA ($m_H$) and mRNA-ribosome complex ($c_H$). Transcriptional repression is modelled by modifying the transcription rate of a gene depending on the concentration of the other protein species. The corresponding ODEs for the expression of a toggle switch are as follows:

$$\dot{m}_{H_i} = \omega_{H_i}(H_{i-1}, e) - \beta_{H_i} \cdot m_{H_i} \cdot R + k^-_{H_i} \cdot c_{H_i} + \Gamma_{H_i}(c_{H_i}, e) - (\lambda + d_m) \cdot m_{H_i}, \tag{S17}$$

$$\dot{c}_{H_i} = \beta_{H_i} \cdot m_{H_i} \cdot R - k^-_{H_i} \cdot c_{H_i} - \Gamma_X(c_{H_i}, e) - \lambda \cdot c_{H_i}, \tag{S18}$$

$$\dot{H}_i = \Gamma_H(c_{H_i}, e) - (\lambda + d_p) \cdot H_i, \tag{S19}$$

where $i \in \{1, 2\}$. The negative regulation function $\omega_{H_i}(H_{i-1}, e)$ regulates the mRNA of one species ($m_{H_i}$) by the other protein ($H_{i-1}$). This itself is affected by the inhibitor which is modelled as affecting the half-saturation constant for repression, $K_H$:

$$\omega_{H_i}(H_{i-1}, K_H, e) = \frac{\alpha_{H_i} \cdot e}{K_{\omega_{H_i}} + e} \cdot \frac{1}{1 + \left(\frac{H_{i-1}}{K_H(I_{i-1})}\right)^{h_H}}, \tag{S20}$$

$$K_H(I_i) = K_H \left(1 + \frac{I_i}{K_I}\right) \tag{S21}$$

We let $H_0 = H_2$ and $I_0 = I_2$ to implement the mutual inhibition between the toggle switch's components. While both inhibitors are represented in these equations, our simulations only explore the effects of one. The additional parameters used are given by:

- Half-saturation constant for repression ($K_H$) = $0.32\,\mathrm{molecs\,cell^{-1}}$;
- Repression Hill coefficient ($h_H$) = 2;
- Half-saturation constant for inhibition ($K_I$) = $1\,\mathrm{molecs\,cell^{-1}}$;
- Length of each synthetic protein ($n_H$) = $300\,\mathrm{aa\,molecs^{-1}}$.

The values of $K_H$, $h_H$ and $K_I$ are those used in the toggle switch simulations from Del Vecchio et al.'s Biomolecular Feedback Systems [12], while the value of $n_H$ is chosen to be equal to the length of all other non-ribosomal proteins (Table S1).

##### S1.4 Adjustments to implement the repressilator

To model a repressilator, each of its three synthetic genes requires ODEs for the protein ($H$), mRNA ($m_H$) and mRNA-ribosome complex ($c_H$). Transcriptional repression is modelled by modifying the transcription rate of a gene depending on the concentration of the another protein species. A parameter is also included for protein degradation rate ($d_p$) as repressilator genes are often designed with degradation tags. The ODEs for the synthetic gene expression are as follows:

$$\dot{m}_{H_i} = \omega_{H_i}(H_{i-1}, e) - \beta_{H_i} \cdot m_{H_i} \cdot R + k^-_{H_i} \cdot c_{H_i} + \Gamma_{H_i}(c_{H_i}, e) - (\lambda + d_m) \cdot m_{H_i}, \tag{S22}$$

$$\dot{c}_{H_i} = \beta_{H_i} \cdot m_{H_i} \cdot R - k^-_{H_i} \cdot c_{H_i} - \Gamma_X(c_{H_i}, e) - \lambda \cdot c_{H_i}, \tag{S23}$$

$$\dot{H}_i = \Gamma_H(c_{H_i}, e) - (\lambda + d_p) \cdot H_i, \tag{S24}$$

where $i \in \{1, 2, 3\}$. The function $\omega_{H_i}(H_{i-1}, e)$ regulates the mRNA of one species ($m_{H_i}$) by the previous protein in the loop ($H_{i-1}$) using negative regulation:

$$\omega_{H_i}(H_{i-1}, e) = \frac{\alpha_{H_i} \cdot e}{K_{\omega_{H_i}} + e} \cdot \frac{1}{1 + \left(\frac{H_{i-1}}{K_H}\right)^{h_H}}. \tag{S25}$$

We let $H_0 = H_3$ to validate the one-way cycle of repression between components. The additional parameters are given by:

- Protein degradation rate ($d_p$) = $4.16\,\mathrm{h}^{-1}$;
- Half-saturation constant for repression ($K_H$) = $100\,\mathrm{molecs\,cell}^{-1}$;
- Hill coefficient for repression ($h_H$) = 2;
- Length of each synthetic protein ($n_H$) = $300\,\mathrm{aa\,molecs}^{-1}$.

The value of $d_p$ is chosen such that oscillations have approximately the same period (3.2 h) as those in the experiments of Elowitz and Leibler [13], while the values of $K_H$ and $h_H$ are chosen to match the values used in Weisse et al.'s [1] simulations, and $n_H$ is chosen to match the size of all non-ribosomal proteins (Table S1).

### S2 Supplementary analysis for model fitting

To demonstrate the potential of our model in capturing real data, we fit simulation results to data from Figure 3 of the fluorescence decay experiments of Sleight et al. [14]. In this study, the authors engineer *E. coli* with variants of the same fluorescent-tagged synthetic construct (‘T9002’ and six variants appended with ‘-A’ to ‘-F’), and record how each population’s fluorescence decays over time due to the onset of mutations. The different designs associated with each construct cause changes to both the mutation probability of the construct and the associated growth rate of the cells. The precise combination of these factors is typically unknown without extensive experimental investigation, however they can be estimated by fitting the key mutation parameters in our model ($\alpha_\mathrm{E}, z_\mathrm{M}, \alpha_\mathrm{I}, z_\mathrm{I}$) to the experimental data.

#### S2.1 Generation and interpretation of parameter values

For each data set, the values of data points were inferred using the online tool ‘WebPlotDigitizer’ (version 4.6). We standardised our values of H-protein production with [14]’s values for fluorescence (measured in $\mathrm{OD}_{600}$) by fixing their maximum-recorded fluorescence value to a simulated $\alpha_\mathrm{E}$ value of $10^4\,\mathrm{molecs\,h^{-1}\,cell^{-1}}$. This required multiplying our simulation values for $H$ by $\frac{\text{highest } \mathrm{OD}_{600} \text{ value recorded by [14]}}{H \text{ in population at } t{=}0\,\mathrm{h} \text{ when } \alpha_\mathrm{E}{=}10^4} = \frac{2.1\times10^4}{9.8\times10^{13}} \approx 2.0\times10^{-10}$. To form a valid comparison to [14]’s data, we converted our ‘time’ unit to ‘generations’ by multiplying the population’s per-cell average growth rate by time, and dividing by $\ln(2)$ to account for ‘doubling time’: generation $= \frac{\sum_{i=1}^{n} \lambda_{X_i}(t)\cdot X_i}{N} \cdot \frac{t}{\ln(2)}$, where $n$ is the number of cell types. The first fraction indicates the population’s average growth rate, where $\lambda_{X_i}(t)$ is the growth rate of cell type $\mathrm{X_i}$ at time $t$, $X_i$ is the number of cells in cell type $\mathrm{X_i}$, and $N$ is the turbidostat’s cell threshold. The fitted curves are shown in Figure S2, with parameter values and fit statistics given in Table S1.

Simulation results were fit to each data set by systematically varying the core mutation parameters ($\alpha_\mathrm{E}, z_\mathrm{M}, \alpha_\mathrm{I}, z_\mathrm{I}$) for both $[s_2, d_1]$ and $[s_3, d_1]$ frameworks until the closest fit was obtained, evaluated by the smallest root mean square deviation. In general, $\alpha_\mathrm{E}$ affects the initial fluorescence value, $z_\mathrm{M}$ affects the length of time over which the overall mutation dynamics occur, while adjusting $\alpha_\mathrm{I}$ and $z_\mathrm{I}$ affects localised regions of the dynamics and in doing so introduces asymmetry in the dynamics. As seen from Table S2, the values of $z_\mathrm{M}$ vary across many orders of magnitude, suggesting that the variation in fluorescence decay can be somewhat attributed to significant differences in mutation probability. Furthermore, the use of $\alpha_\mathrm{I}$ and $z_\mathrm{I}$ for all but

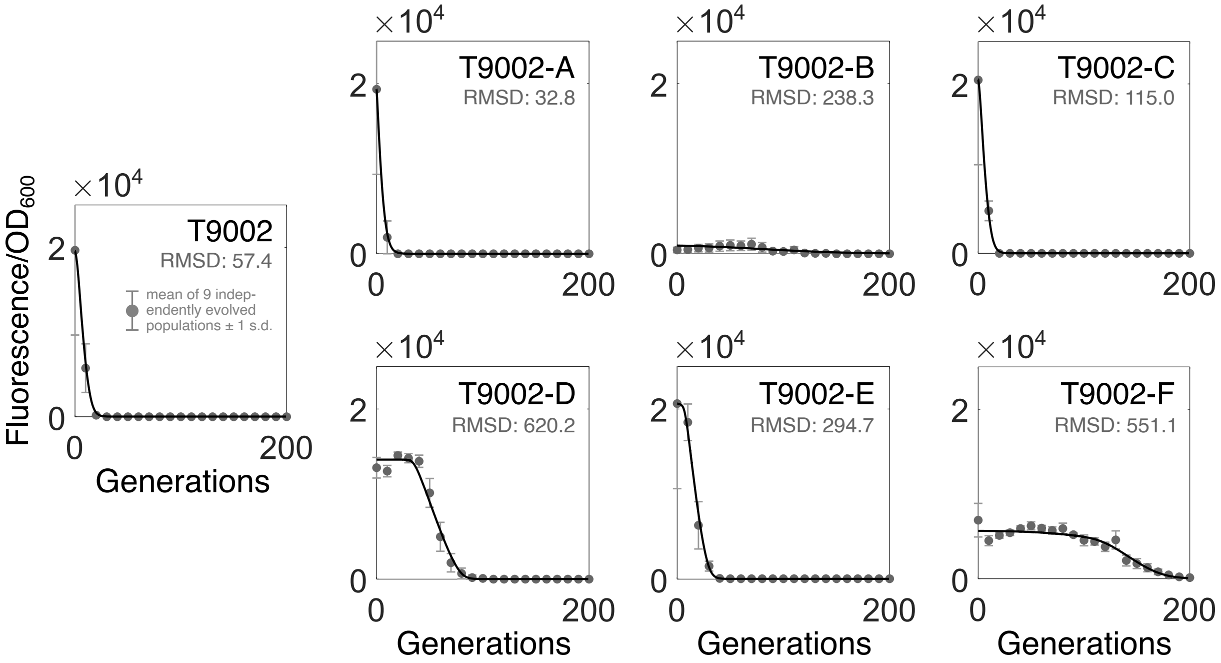


**Figure S2: Fitting simulation results to experimental data from [14].** Our $[s_3, d_1]$ framework is used to fit simulations to [14]'s fluorescence decay experiments of the synthetic construct T9002 and its six variants (appended with -A to -F). Dots represent the mean of nine experimental repeats, with error bars representing one standard deviation. Simulation data is given as a black line. The unit of 'time' from our models is converted to 'generations' as explained in the main text of Section S2. The parameters used for each fit are given in Table S2. RMSD: root mean square deviation.

**Table S2: Parameter values used to fit simulations to experimental data from [14].** Four parameters ($\alpha_{\mathrm{E}}, z_{\mathrm{M}}, \alpha_{\mathrm{I}}, z_{\mathrm{I}}$) are adjusted to best approximate the data. RMSD: root mean square deviation. 'x': parameter is not used in the fit. The 'relative instability predictor' (RIP) score denotes the relative mutation rate of a sequence calculated by [3]'s EFM calculator. This is used to convert a construct's baseline mutation rate to a RIP-adjusted mutation rate as 'adjusted = baseline · RIP score'.
**a**: $1-(1-\mu)^n$, where $\mu = 4.1\times10^{-10}\,\mathrm{nt}^{-1}\,\mathrm{gen}^{-1}$ [15] and $n$ is the number of nucleotides in a construct. For the values to be compatible with our model, we treat rates as probabilities, which is a good approximation when these values are small (see the text of Section S2).

| Construct | Best fit parameters $[\alpha_{\mathrm{E}}, z_{\mathrm{M}}, \alpha_{\mathrm{I}}, z_{\mathrm{I}}]$ | RMSD | Baseline mutation probability[a] | RIP score | RIP-adjusted mutation probability |
|---|---|---|---|---|---|
| T9002 | $[9.0\times10^{3}, 2.0\times10^{-2}, 5.0\times10^{3}, 4.0\times10^{-2}]$ | 57.4 | $7.97\times10^{-7}$ | 486.3 | $3.88\times10^{-4}$ |
| T9002-A | $[8.8\times10^{3}, 1.0\times10^{-1}, 2.5\times10^{3}, 1.0\times10^{-1}]$ | 32.8 | $7.97\times10^{-7}$ | 100.8 | $8.04\times10^{-5}$ |
| T9002-B | $[2.0\times10^{2}, 4.0\times10^{-3}$, x, x$]$ | 238.3 | $7.97\times10^{-7}$ | 322.1 | $2.57\times10^{-4}$ |
| T9002-C | $[9.9\times10^{3}, 4.0\times10^{-2}, 2.8\times10^{3}, 2.0\times10^{-2}]$ | 115.0 | $7.84\times10^{-7}$ | 235.3 | $1.84\times10^{-4}$ |
| T9002-D | $[5.0\times10^{3}, 2.2\times10^{-10}$, x, x$]$ | 620.2 | $7.84\times10^{-7}$ | 102.6 | $8.04\times10^{-5}$ |
| T9002-E | $[1.0\times10^{4}, 7.0\times10^{-5}, 4.1\times10^{3}, 3.0\times10^{-3}]$ | 294.7 | $7.91\times10^{-7}$ | 101.6 | $8.04\times10^{-5}$ |
| T9002-F | $[1.4\times10^{3}, 4.0\times10^{-10}, 1.0\times10^{3}, 1.0\times10^{-3}]$ | 551.1 | $7.63\times10^{-7}$ | 182.9 | $1.40\times10^{-4}$ |

two of the constructs (T9002-B and T9002-D) to achieve an optimal fit suggest the presence of mutation heterogeneity within the experiments.

In order to evaluate the accuracy of our parameter values, extensive sequencing data would be required to confirm the distribution of mutations and their frequencies within each experiment. The results of [14] do not include this, however we can compare our parameter for $z_{\mathrm{M}}$ to existing tools that estimate the mutation probability of a given DNA sequence, such as

Jack et al.'s EFM calculator [3]. This tool generates a 'relative instability predictor' (RIP) score from an input sequence which suggests how mutagenic it is relative to the baseline mutation rate. By taking *E. coli*'s baseline as $4.1{\times}10^{-10}\,\mathrm{nt}^{-1}\,\mathrm{gen}^{-1}$ [15], a RIP-adjusted mutation rate can be calculated by multiplying the baseline rate by the RIP score. Given our framework requires values as 'probabilities', we can calculate these from 'rate' values using the Poisson distribution: this gives a probability ($p$) of X events happening if they occur at some time-averaged rate, $\Lambda$: $p(X = k) = \frac{\Lambda^k \cdot e^{-\Lambda}}{k!}$. For the probability of a mutation occurring in one generation, $p(X = 1) = \frac{\Lambda^1 \cdot e^{-\Lambda}}{1!} = \Lambda \cdot e^{-\Lambda}$. When values of $\Lambda$ are small, $\Lambda \cdot e^{-\Lambda} \approx \Lambda$, meaning we can use 'probability $\approx$ rate' as a reasonable assumption and directly compare values of the RIP-adjusted mutation rate with our parameter for mutation probability.

By comparing the fitted values of $z_{\mathrm{M}}$ to those derived from the EFM calculator, we see that there is varying agreement between constructs. T9002-B and T9002-E have similar pairs of values ($[z_{\mathrm{M}}, \mathrm{RIP}] = [3.0{\times}10^{-3}, 2.6{\times}10^{-4}]$ and $[z_{\mathrm{M}}, \mathrm{RIP}] = [8.0{\times}10^{-5}, 8.0{\times}10^{-5}]$, respectively), while T9002-D and T9002-F have large deviations ($[z_{\mathrm{M}}, \mathrm{RIP}] = [2.2{\times}10^{-10}, 8.0{\times}10^{-5}]$ and $[z_{\mathrm{M}}, \mathrm{RIP}] = [5.0{\times}10^{-10}, 1.4{\times}10^{-4}]$, respectively). In their paper, Jack et al. find a correlation between the evolutionary half-lives of T9002 and T9002-E and a change in RIP scores, however they do not repeat the analysis for the other T9002 variants, some of which have significantly higher evolutionary half-lives (for example T9002-B and T9002-F). When applying the EFM calculator to all T9002 constructs, we find that the range of values for RIP scores and mutation probabilities are within a single order of magnitude (Table S2), and based on our modelling this is insufficient to capture the range of mutation dynamics observed, even when freely adjusting $\alpha_{\mathrm{E}}$, $\alpha_{\mathrm{I}}$ and $z_{\mathrm{I}}$. This suggests that, in some settings, the mutation dynamics in a system are likely more complex than what the EFM calculator can account for. As shown by our analysis, this could be corrected by allowing a larger variation in the parameters associated with mutation probability.

#### S2.2 Generation of RIP scores

We calculated RIP scores of the T9002 constructs using version 1.0.1 of the EFM calculator. The sequences used as input were derived by searching for each part that Sleight et al. report in their study within iGEM's 'Registry of Standard Biological Parts' and joining them together in the correct order and orientation. The DNA sequences we inferred for the individual genetic parts and for the T9002 constructs are provided below.

**B0010**: ccaggcatcaaataaaacgaaaggctcagtcgaaagactgggcctttcgttttatctgttgtttgtcggtg
aacgctctc **B0011**: agagaatataaaaagccagattattaatccggcttttttattattt **B0012**: tcacactg
gctcaccttcgggtgggcctttctgcgtttata **J61048**: ccggcttatcggtcagtttcacctgatttacgtaaaa
acccgcttcggcgggtttttgcttttggaggggcagaaagatgaatgactgtccacgacgctatacccaaaagaaa

**T9002**: tccctatcagtgatagagattgacatccctatcagtgatagagatactgagcactactagagaaagagga
gaaatactagatgaaaaacataaatgccgacgacacatacagaataattaataaaattaaagcttgtagaagcaataat
gatattaatcaatgcttatctgatatgactaaaatggtacattgtgaatattatttactcgcgatcatttatcctcatt
ctatggttaaatctgatatttcaatcctagataattaccctaaaaaatggaggcaatattatgatgacgctaatttaat
aaaatatgatcctatagtagattattctaactccaatcattcaccaattaattggaatatatttgaaaacaatgctgta
aataaaaaatctccaaatgtaattaaagaagcgaaaacatcaggtcttatcactgggtttagtttccctattcatacgg
ctaacaatggcttcggaatgcttagttttgcacattcagaaaaagacaactatatagatagtttatttttacatgcgtg
tatgaacataccattaattgttccttctctagttgataattatcgaaaaataaatatagcaaataataaatcaaacaac
gatttaaccaaaagagaaaaagaatgtttagcgtgggcatgcgaaggaaaaagctcttgggatatttcaaaaatattag
gttgcagtgagcgtactgtcactttccatttaaccaatgcgcaaatgaaactcaatacaacaaaccgctgccaaagtat
ttctaaagcaattttaacaggagcaattgattgcccatactttaaaaattaataacactgatagtgctagtgtagatca
ctactagagccaggcatcaaataaaacgaaaggctcagtcgaaagactgggcctttcgttttatctgttgtttgtcggt
gaacgctctctactagagtcacactggctcaccttcgggtgggcctttctgcgtttatatactagagacctgtaggatc
gtacaggtttacgcaagaaaatggtttgttatagtcgaataaatactagagtcacacaggaaagtactagatgcgtaaa
ggagaagaacttttcactggagttgtcccaattcttgttgaattagatggtgatgttaatgggcacaaattttctgtca
gtggagagggtgaaggtgatgcaacatacggaaaacttacccttaaatttatttgcactactggaaaactacctgttcc
atggccaacacttgtcactactttcggttatggtgttcaatgctttgcgagatacccagatcatatgaaacagcatgac
tttttcaagagtgccatgcccgaaggttatgtacaggaaagaactatatttttcaaagatgacgggaactacaagacac
gtgctgaagtcaagtttgaaggtgatacccttgttaatagaatcgagttaaaaggtattgattttaaagaagatggaaa
cattcttggacacaaattggaatacaactataactcacacaatgtatacatcatggcagacaaacaaaagaatggaatc
aaagttaacttcaaaattagacacaacattgaagatggaagcgttcaactagcagaccattatcaacaaaatactccaa
ttggcgatggccctgtccttttaccagacaaccattacctgtccacacaatctgccctttcgaaagatcccaacgaaaa
gagagaccacatggtccttcttgagtttgtaacagctgctgggattacacatggcatggatgaactatacaaataataa
tactagagccaggcatcaaataaaacgaaaggctcagtcgaaagactgggcctttcgttttatctgttgtttgtcggtg
aacgctctctactagagtcacactggctcaccttcgggtgggcctttctgcgtttata

**T9002-A**: tccctatcagtgatagagattgacatccctatcagtgatagagatactgagcactactagagaaagag
gagaaatactagatgaaaaacataaatgccgacgacacatacagaataattaataaaattaaagcttgtagaagcaata
atgatattaatcaatgcttatctgatatgactaaaatggtacattgtgaatattatttactcgcgatcatttatcctca
ttctatggttaaatctgatatttcaatcctagataattaccctaaaaaatggaggcaatattatgatgacgctaattta
ataaaatatgatcctatagtagattattctaactccaatcattcaccaattaattggaatatatttgaaaacaatgctg
taaataaaaaatctccaaatgtaattaaagaagcgaaaacatcaggtcttatcactgggtttagtttccctattcatac
ggctaacaatggcttcggaatgcttagttttgcacattcagaaaaagacaactatatagatagtttatttttacatgcg
tgtatgaacataccattaattgttccttctctagttgataattatcgaaaaataaatatagcaaataataaatcaaaca
acgatttaaccaaaagagaaaaagaatgtttagcgtgggcatgcgaaggaaaaagctcttgggatatttcaaaaatatt
aggttgcagtgagcgtactgtcactttccatttaaccaatgcgcaaatgaaactcaatacaacaaaccgctgccaaagt
atttctaaagcaattttaacaggagcaattgattgcccatactttaaaaattaataacactgatagtgctagtgtagat
cactactagagccaggcatcaaataaaacgaaaggctcagtcgaaagactgggcctttcgttttatctgttgtttgtcg
gtgaacgctctctactagagtcacactggctcaccttcgggtgggcctttctgcgtttatatactagagacctgtagga
tcgtacaggtttacgcaagaaaatggtttgttatagtcgaataaatactagagtcacacaggaaagtactagatgcgta
aaggagaagaacttttcactggagttgtcccaattcttgttgaattagatggtgatgttaatgggcacaaattttctgt
cagtggagagggtgaaggtgatgcaacatacggaaaacttacccttaaatttatttgcactactggaaaactacctgtt
ccatggccaacacttgtcactactttcggttatggtgttcaatgctttgcgagatacccagatcatatgaaacagcatg
actttttcaagagtgccatgcccgaaggttatgtacaggaaagaactatatttttcaaagatgacgggaactacaagac
acgtgctgaagtcaagtttgaaggtgatacccttgttaatagaatcgagttaaaaggtattgattttaaagaagatgga
aacattcttggacacaaattggaatacaactataactcacacaatgtatacatcatggcagacaaacaaaagaatggaa

tcaaagttaacttcaaaattagacacaacattgaagatggaagcgttcaactagcagaccattatcaacaaaatactcc
aattggcgatggccctgtccttttaccagacaaccattacctgtccacacaatctgccctttcgaaagatcccaacgaa
aagagagaccacatggtccttcttgagtttgtaacagctgctgggattacacatggcatggatgaactatacaaataat
aatactagagtataaacgcagaaaggcccacccgaaggtgagccagtgtgatactagaggagagcgttcaccgacaaac
aacagataaaacgaaaggcccagtctttcgactgagcctttcgttttatttgatgcctgg

**T9002-B**: tccctatcagtgatagagattgacatccctatcagtgatagagatactgagcactactagagaaagag
gagaaatactagatgaaaaacataaatgccgacgacacatacagaataattaataaaattaaagcttgtagaagcaata
atgatattaatcaatgcttatctgatatgactaaaatggtacattgtgaatattatttactcgcgatcatttatcctca
ttctatggttaaatctgatatttcaatcctagataattaccctaaaaaatggaggcaatattatgatgacgctaattta
ataaaatatgatcctatagtagattattctaactccaatcattcaccaattaattggaatatatttgaaaacaatgctg
taaataaaaaatctccaaatgtaattaaagaagcgaaaacatcaggtcttatcactgggtttagtttccctattcatac
ggctaacaatggcttcggaatgcttagttttgcacattcagaaaaagacaactatatagatagtttatttttacatgcg
tgtatgaacataccattaattgttccttctctagttgataattatcgaaaaataaatatagcaaataataaatcaaaca
acgatttaaccaaaagagaaaaagaatgtttagcgtgggcatgcgaaggaaaaagctcttgggatatttcaaaaatatt
aggttgcagtgagcgtactgtcactttccatttaaccaatgcgcaaatgaaactcaatacaacaaaccgctgccaaagt
atttctaaagcaattttaacaggagcaattgattgcccatactttaaaaattaataacactgatagtgctagtgtagat
cactactagagccaggcatcaaataaaacgaaaggctcagtcgaaagactgggcctttcgttttatctgttgtttgtcg
gtgaacgctctctactagagtcacactggctcaccttcgggtgggcctttctgcgtttatatactagagacctgtagga
tcgtacaggtttacgcaagaaaatggtttgttatagtcgaataaatactagagtcacacaggaaagtactagatgcgta
aaggagaagaacttttcactggagttgtcccaattcttgttgaattagatggtgatgttaatgggcacaaattttctgt
cagtggagagggtgaaggtgatgcaacatacggaaaacttacccttaaatttatttgcactactggaaaactacctgtt
ccatggccaacacttgtcactactttcggttatggtgttcaatgctttgcgagatacccagatcatatgaaacagcatg
actttttcaagagtgccatgcccgaaggttatgtacaggaaagaactatatttttcaaagatgacgggaactacaagac
acgtgctgaagtcaagtttgaaggtgatacccttgttaatagaatcgagttaaaaggtattgattttaaagaagatgga
aacattcttggacacaaattggaatacaactataactcacacaatgtatacatcatggcagacaaacaaaagaatggaa
tcaaagttaacttcaaaattagacacaacattgaagatggaagcgttcaactagcagaccattatcaacaaaatactcc
aattggcgatggccctgtccttttaccagacaaccattacctgtccacacaatctgccctttcgaaagatcccaacgaa
aagagagaccacatggtccttcttgagtttgtaacagctgctgggattacacatggcatggatgaactatacaaataat
aatactagagtcacactggctcaccttcgggtgggcctttctgcgtttatatactagagccaggcatcaaataaaacga
aaggctcagtcgaaagactgggcctttcgttttatctgttgtttgtcggtgaacgctctc

**T9002-C**: tccctatcagtgatagagattgacatccctatcagtgatagagatactgagcactactagagaaagag
gagaaatactagatgaaaaacataaatgccgacgacacatacagaataattaataaaattaaagcttgtagaagcaata
atgatattaatcaatgcttatctgatatgactaaaatggtacattgtgaatattatttactcgcgatcatttatcctca
ttctatggttaaatctgatatttcaatcctagataattaccctaaaaaatggaggcaatattatgatgacgctaattta
ataaaatatgatcctatagtagattattctaactccaatcattcaccaattaattggaatatatttgaaaacaatgctg
taaataaaaaatctccaaatgtaattaaagaagcgaaaacatcaggtcttatcactgggtttagtttccctattcatac
ggctaacaatggcttcggaatgcttagttttgcacattcagaaaaagacaactatatagatagtttatttttacatgcg
tgtatgaacataccattaattgttccttctctagttgataattatcgaaaaataaatatagcaaataataaatcaaaca
acgatttaaccaaaagagaaaaagaatgtttagcgtgggcatgcgaaggaaaaagctcttgggatatttcaaaaatatt
aggttgcagtgagcgtactgtcactttccatttaaccaatgcgcaaatgaaactcaatacaacaaaccgctgccaaagt
atttctaaagcaattttaacaggagcaattgattgcccatactttaaaaattaataacactgatagtgctagtgtagat
cactactagagccaggcatcaaataaaacgaaaggctcagtcgaaagactgggcctttcgttttatctgttgtttgtcg
gtgaacgctctctactagagtcacactggctcaccttcgggtgggcctttctgcgtttatatactagagacctgtagga
tcgtacaggtttacgcaagaaaatggtttgttatagtcgaataaatactagagtcacacaggaaagtactagatgcgta
aaggagaagaacttttcactggagttgtcccaattcttgttgaattagatggtgatgttaatgggcacaaattttctgt
cagtggagagggtgaaggtgatgcaacatacggaaaacttacccttaaatttatttgcactactggaaaactacctgtt
ccatggccaacacttgtcactactttcggttatggtgttcaatgctttgcgagatacccagatcatatgaaacagcatg
actttttcaagagtgccatgcccgaaggttatgtacaggaaagaactatatttttcaaagatgacgggaactacaagac
acgtgctgaagtcaagtttgaaggtgatacccttgttaatagaatcgagttaaaaggtattgattttaaagaagatgga

aacattcttggacacaaattggaatacaactataactcacacaatgtatacatcatggcagacaaacaaaagaatggaa
tcaaagttaacttcaaaattagacacaacattgaagatggaagcgttcaactagcagaccattatcaacaaaatactcc
aattggcgatggccctgtccttttaccagacaaccattacctgtccacacaatctgccctttcgaaagatcccaacgaa
aagagagaccacatggtccttcttgagtttgtaacagctgctgggattacacatggcatggatgaactatacaaataat
aatactagagtcacactggctcaccttcgggtgggcctttctgcgtttatatactagagagagaatataaaaagccaga
ttattaatccggctttttttattattt

**T9002-D**: tccctatcagtgatagagattgacatccctatcagtgatagagatactgagcactactagagaaagag
gagaaatactagatgaaaaacataaatgccgacgacacatacagaataattaataaaattaaagcttgtagaagcaata
atgatattaatcaatgcttatctgatatgactaaaatggtacattgtgaatattatttactcgcgatcatttatcctca
ttctatggttaaatctgatatttcaatcctagataattaccctaaaaaatggaggcaatattatgatgacgctaattta
ataaaatatgatcctatagtagattattctaactccaatcattcaccaattaattggaatatatttgaaaacaatgctg
taaataaaaaatctccaaatgtaattaaagaagcgaaaacatcaggtcttatcactgggtttagtttccctattcatac
ggctaacaatggcttcggaatgcttagttttgcacattcagaaaaagacaactatatagatagtttatttttacatgcg
tgtatgaacataccattaattgttccttctctagttgataattatcgaaaaataaatatagcaaataataaatcaaaca
acgatttaaccaaaagagaaaaagaatgtttagcgtgggcatgcgaaggaaaaagctcttgggatatttcaaaaatatt
aggttgcagtgagcgtactgtcactttccatttaaccaatgcgcaaatgaaactcaatacaacaaaccgctgccaaagt
atttctaaagcaattttaacaggagcaattgattgcccatactttaaaaattaataacactgatagtgctagtgtagat
cactactagagccaggcatcaaataaaacgaaaggctcagtcgaaagactgggcctttcgttttatctgttgtttgtcg
gtgaacgctctctactagagtcacactggctcaccttcgggtgggcctttctgcgtttatatactagagacctgtagga
tcgtacaggtttacgcaagaaaatggtttgttatagtcgaataaatactagagtcacacaggaaagtactagatgcgta
aaggagaagaacttttcactggagttgtcccaattcttgttgaattagatggtgatgttaatgggcacaaattttctgt
cagtggagagggtgaaggtgatgcaacatacggaaaacttacccttaaatttatttgcactactggaaaactacctgtt
ccatggccaacacttgtcactactttcggttatggtgttcaatgctttgcgagatacccagatcatatgaaacagcatg
actttttcaagagtgccatgcccgaaggttatgtacaggaaagaactatatttttcaaagatgacgggaactacaagac
acgtgctgaagtcaagtttgaaggtgatacccttgttaatagaatcgagttaaaaggtattgattttaaagaagatgga
aacattcttggacacaaattggaatacaactataactcacacaatgtatacatcatggcagacaaacaaaagaatggaa
tcaaagttaacttcaaaattagacacaacattgaagatggaagcgttcaactagcagaccattatcaacaaaatactcc
aattggcgatggccctgtccttttaccagacaaccattacctgtccacacaatctgccctttcgaaagatcccaacgaa
aagagagaccacatggtccttcttgagtttgtaacagctgctgggattacacatggcatggatgaactatacaaataat
aatactagagaaataataaaaaagccggattaataatctggctttttatattctcttactagagtataaacgcagaaag
gcccacccgaaggtgagccagtgtga

**T9002-E**: tccctatcagtgatagagattgacatccctatcagtgatagagatactgagcactactagagaaagag
gagaaatactagatgaaaaacataaatgccgacgacacatacagaataattaataaaattaaagcttgtagaagcaata
atgatattaatcaatgcttatctgatatgactaaaatggtacattgtgaatattatttactcgcgatcatttatcctca
ttctatggttaaatctgatatttcaatcctagataattaccctaaaaaatggaggcaatattatgatgacgctaattta
ataaaatatgatcctatagtagattattctaactccaatcattcaccaattaattggaatatatttgaaaacaatgctg
taaataaaaaatctccaaatgtaattaaagaagcgaaaacatcaggtcttatcactgggtttagtttccctattcatac
ggctaacaatggcttcggaatgcttagttttgcacattcagaaaaagacaactatatagatagtttatttttacatgcg
tgtatgaacataccattaattgttccttctctagttgataattatcgaaaaataaatatagcaaataataaatcaaaca
acgatttaaccaaaagagaaaaagaatgtttagcgtgggcatgcgaaggaaaaagctcttgggatatttcaaaaatatt
aggttgcagtgagcgtactgtcactttccatttaaccaatgcgcaaatgaaactcaatacaacaaaccgctgccaaagt
atttctaaagcaattttaacaggagcaattgattgcccatactttaaaaattaataacactgatagtgctagtgtagat
cactactagagccaggcatcaaataaaacgaaaggctcagtcgaaagactgggcctttcgttttatctgttgtttgtcg
gtgaacgctctctactagagtcacactggctcaccttcgggtgggcctttctgcgtttatatactagagacctgtagga
tcgtacaggtttacgcaagaaaatggtttgttatagtcgaataaatactagagtcacacaggaaagtactagatgcgta
aaggagaagaacttttcactggagttgtcccaattcttgttgaattagatggtgatgttaatgggcacaaattttctgt
cagtggagagggtgaaggtgatgcaacatacggaaaacttacccttaaatttatttgcactactggaaaactacctgtt
ccatggccaacacttgtcactactttcggttatggtgttcaatgctttgcgagatacccagatcatatgaaacagcatg
actttttcaagagtgccatgcccgaaggttatgtacaggaaagaactatatttttcaaagatgacgggaactacaagac

acgtgctgaagtcaagtttgaaggtgataccctgttaatagaatcgagttaaaaggtattgattttaaagaagatgga
aacattcttggacacaaattggaatacaactataactcacacaatgtatacatcatggcagacaaacaaaagaatggaa
tcaaagttaacttcaaaattagacacaacattgaagatggaagcgttcaactagcagaccattatcaacaaaatactcc
aattggcgatggccctgtccttttaccagacaaccattacctgtccacacaatctgccctttcgaaagatcccaacgaa
aagagagaccacatggtccttcttgagtttgtaacagctgctgggattacacatggcatggatgaactatacaaataat
aatactagagccggcttatcggtcagtttcacctgatttacgtaaaaacccgcttcggcgggtttttgcttttggaggg
gcagaaagatgaatgactgtccacgacgctatacccaaaagaaa

**T9002-F**: tccctatcagtgatagagattgacatccctatcagtgatagagatactgagcactactagagaaagag
gagaaatactagatgaaaaacataaatgccgacgacacatacagaataattaataaaattaaagcttgtagaagcaata
atgatattaatcaatgcttatctgatatgactaaaatggtacattgtgaatattatttactcgcgatcatttatcctca
ttctatggttaaatctgatatttcaatcctagataattaccctaaaaaatggaggcaatattatgatgacgctaattta
ataaaatatgatcctatagtagattattctaactccaatcattcaccaattaattggaatatatttgaaaacaatgctg
taaataaaaaatctccaaatgtaattaaagaagcgaaaacatcaggtcttatcactgggtttagtttccctattcatac
ggctaacaatggcttcggaatgcttagttttgcacattcagaaaaagacaactatatagatagtttatttttacatgcg
tgtatgaacataccattaattgttccttctctagttgataattatcgaaaaataaatatagcaaataataaatcaaaca
acgatttaaccaaaagagaaaaagaatgtttagcgtgggcatgcgaaggaaaaagctcttgggatatttcaaaaatatt
aggttgcagtgagcgtactgtcactttccatttaaccaatgcgcaaatgaaactcaatacaacaaaccgctgccaaagt
atttctaaagcaattttaacaggagcaattgattgcccatactttaaaaattaataacactgatagtgctagtgtagat
cactactagagccaggcatcaaataaaacgaaaggctcagtcgaaagactgggcctttcgttttatctgttgtttgtcg
gtgaacgctctctactagagtcacactggctcaccttcgggtgggcctttctgcgtttatatactagagacctgtagga
tcgtacaggtttacgcaagaaaatggtttgttatagtcgaataaatactagagtcacacaggaaagtactagatgcgta
aaggagaagaacttttcactggagttgtcccaattcttgttgaattagatggtgatgttaatgggcacaaattttctgt
cagtggagagggtgaaggtgatgcaacatacggaaaacttacccttaaatttatttgcactactggaaaactacctgtt
ccatggccaacacttgtcactactttcggttatggtgttcaatgctttgcgagatacccagatcatatgaaacagcatg
actttttcaagagtgccatgcccgaaggttatgtacaggaaagaactatatttttcaaagatgacgggaactacaagac
acgtgctgaagtcaagtttgaaggtgataccctgttaatagaatcgagttaaaaggtattgattttaaagaagatgga
aacattcttggacacaaattggaatacaactataactcacacaatgtatacatcatggcagacaaacaaaagaatggaa
tcaaagttaacttcaaaattagacacaacattgaagatggaagcgttcaactagcagaccattatcaacaaaatactcc
aattggcgatggccctgtccttttaccagacaaccattacctgtccacacaatctgccctttcgaaagatcccaacgaa
aagagagaccacatggtccttcttgagtttgtaacagctgctgggattacacatggcatggatgaactatacaaataat
aatactagagagagaatataaaaagccagattattaatccggctttttattattt

### S3 Supplementary simulations for gene regulatory networks

#### S3.1 State-wise dynamics for the toggle switch

To gain a more detailed understanding of how the mutation dynamics evolve in the toggle switch simulations from Section 4 in the main text, we can plot the number of cells in each state over time. As before, these simulations start with all cells in the E-state with steady state expression of protein-A and no expression of protein-B. The inhibitor of protein-A ($I_A$) is not added in these cases.

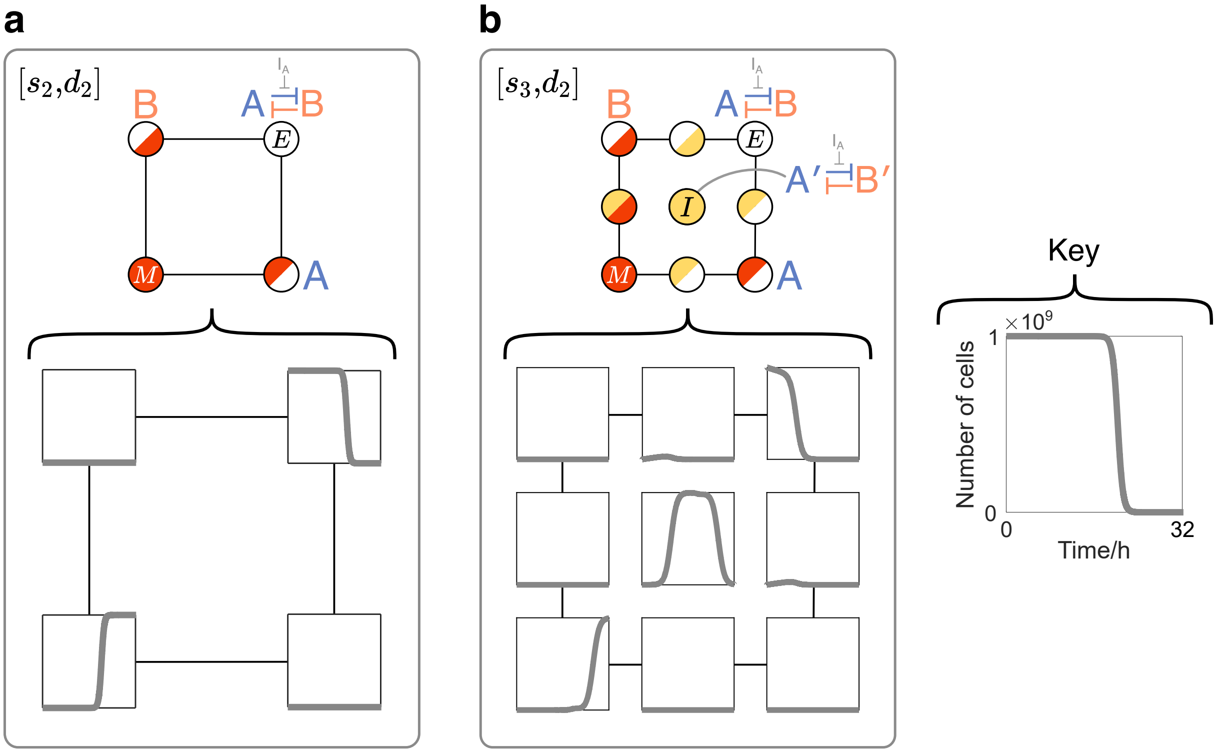


**Figure S3: Additional data for the toggle switch simulations.** **(a)** The number of cells in each state for the $[s_2, d_2]$ framework. Each plot corresponds to the number of cells within the matching mutation state above it. For each synthetic gene, $[\alpha, z] = [10^5 \text{ molecs h}^{-1} \text{ cell}^{-1}, 10^{-6}]$. **(b)** As in (a), except for the $[s_3, d_2]$ framework. For each synthetic gene, $[\alpha_\text{E}, z_\text{M}, \alpha_\text{E}, z_\text{I}] = [10^5 \text{ molecs h}^{-1} \text{ cell}^{-1}, 10^{-6}, 5{\times}10^3 \text{ molecs h}^{-1} \text{ cell}^{-1}, 5{\times}10^{-2}]$.

In Figure S3, each square plot corresponds to the number of cells within the matching mutation state above it, such that the top-right plot corresponds with the top-right mutation state, and so on. For the $[s_2, d_2]$ framework, the most significant changes in cell number occur to the E-state (top-right) and M-state (bottom-left). For the $[s_3, d_2]$ framework, the most significant changes occur to the E-state, I-state and M-state. States with no visible change in cell number still undergo dynamics but to a much smaller degree and so are not visible when using equivalent y-axis scales.

#### S3.2 State-wise dynamics for the repressilator

To gain a more detailed understanding of how the mutation dynamics evolve in the repressilator simulations from Section 5 in the main text, we can plot the number of cells and the average protein concentrations per cell in each state over time. As before, these simulations start with all cells in the E-state and with uniform protein oscillations.

In Figure S4b, we apply the $[s_2, d_3]$ framework such that only fully-inactivating promoter mutations are permitted. From the plots of number of cells in each state, it can be seen that

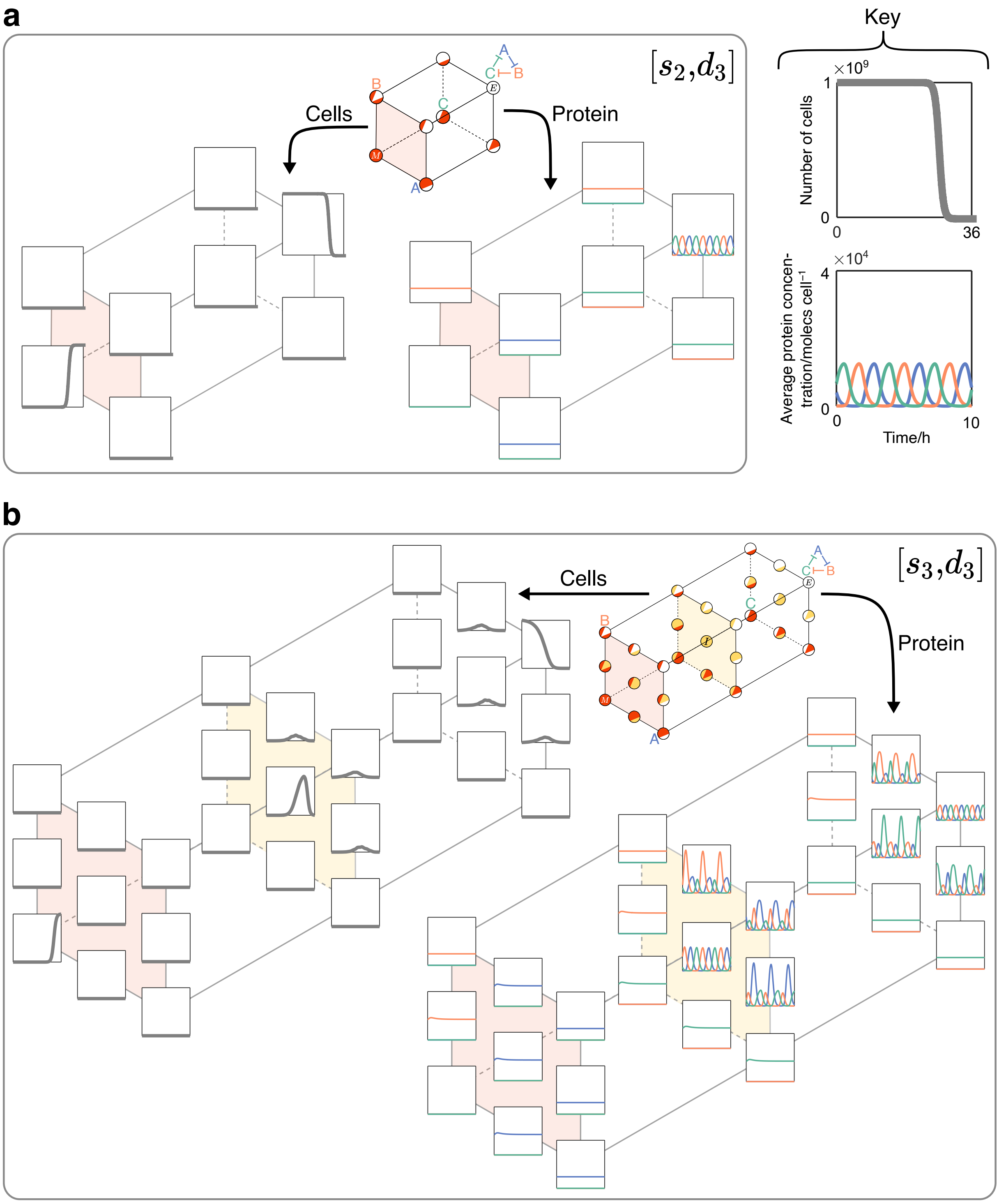


**Figure S4: Additional data for the repressilator simulations.** **(a)** The number of cells (left-hand graphs) and the average concentration of each synthetic protein per cell (right-hand graphs) in each state using the $[s_2, d_2]$ framework. Each plot corresponds to the matching mutation state in the displayed framework. For each synthetic gene, $[\alpha, z] = [10^5$ molecs h$^{-1}$ cell$^{-1}$, $10^{-6}]$. **(c)** As in (b), but additionally modelling partially-inactivating mutations to each synthetic gene's promoter using the $[s_3, d_3]$ framework. For each synthetic gene, $[\alpha_\mathrm{E}, z_\mathrm{M}, \alpha_\mathrm{E}, z_\mathrm{I}] = [10^5$ molecs h$^{-1}$ cell$^{-1}$, $10^{-6}$, $1.2{\times}10^4$ molecs h$^{-1}$ cell$^{-1}$, $1.2{\times}10^{-2}]$.

the most significant changes occur to the E-state (top-right) and M-state (bottom-left). As with the toggle switch plots, states with no visible change in cell number still undergo dynamics but to a much smaller degree and so are not visible when using equivalent y-axis scales. For the concentrations of synthetic proteins, it can be seen how disabling any one gene removes the oscillating behaviour and permits the non-repressed gene to be expressed at a constant value.

A similar analysis can be performed for the $[s_3, d_3]$ framework. As noted in Section 5.2 in the main text, permitting partially-inactivating mutations to a repressilator gives a greater selective advantage for cells in intermediate states to persist in the turbidostat over extended periods of time, owing to their higher growth rate. These effects are reflected in the varied dynamics within other states with partially-inactivating mutations, such as those in the 'upper-right' portion of the framework. The corresponding synthetic protein quantities of these states produce varied oscillation patterns, owing to the fact that all three genes are still active to some degree but with varied repression effects. Fully disabling one gene causes a loss in oscillations, as seen for the states in the 'lower-left' portion of the framework.

### S4 Supplementary mutation modelling

In our analyses so far, we have focused on mutations that decrease the activity of a part and have explored the burden-driven consequences of these. It is conceivable, however, that mutations could increase a part's activity, such as is the goal of many directed evolution experiments. Our framework can easily consider such mutations by simply specifying that a part has a higher activity than in the E-state when a partial mutation occurs. This is achieved by modifying the appropriate value within the input vector '$[p]$' (Figure S1). Using this idea, frameworks can now be represented by placing function-decreasing states downstream of the E-state and function-increasing states upstream of the E-state, with the latter type being distinguished using blue shading and the symbol 'M*'.

For the new framework structures shown in Figure S5, the total number of states in each is the same as in our previously-depicted $s_3$ frameworks. Despite this, as it is now possible to transition from a state with lower activity to one with higher activity, it introduces the question as to whether reverse mutations should be modelled. For example, after transitioning from the E-state to the M-state, could we then transition from the M-state to the M*-state, or even back to the E-state again? And if so, what would the probability of such transitions be? Answering

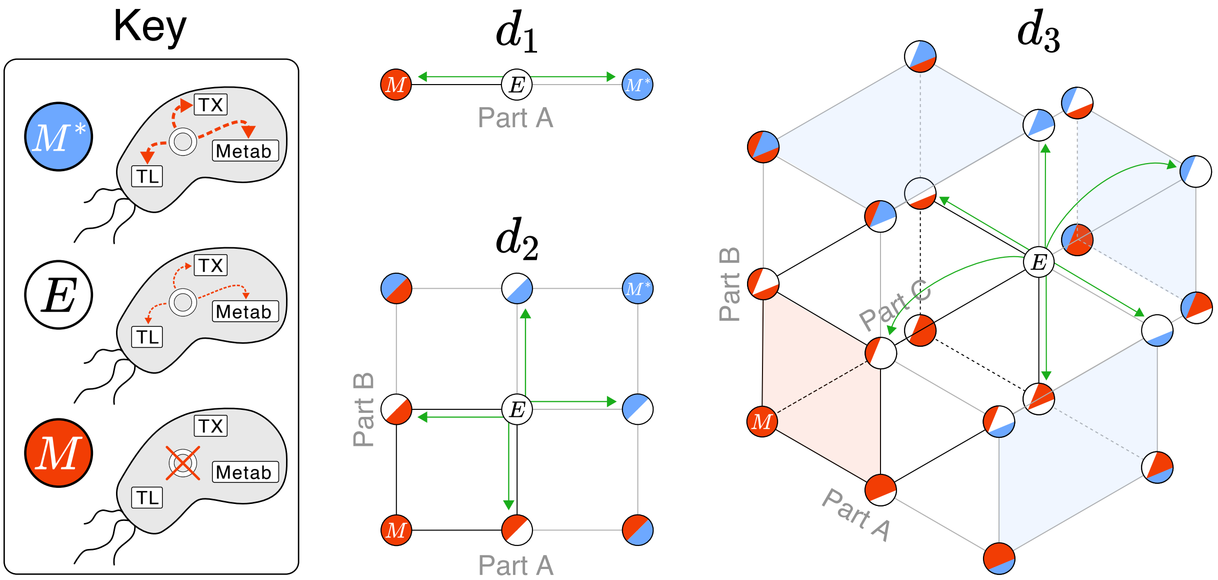


**Figure S5: Framework structures for mutations that both decrease and increase the activity of synthetic parts.** States that depict cells with a function-decreasing mutation are coloured in red and positioned downstream of the E-state, while those with a function-increasing mutation are coloured in blue and positioned upstream of the E-state. Frameworks for the first three dimensions are shown. Some states in the $d_3$ framework are removed for clarity. Some areas between states are shaded to help visual clarity.

such questions with any reliability would require a lot of extensive knowledge about a construct's sequence and the potential for that sequence to change in many different ways. Such approaches may require implementing dynamic evolutionary landscapes, such as those discussed by Castle et al. [16].

Regardless of how transitions are implemented, the effect of function-increasing mutations on the resulting dynamics will likely be negligible. This is due to the phenomenon explored with the repressilator during Section 5 in the main text: any mutations that increase the burdensome effects of synthetic gene expression will decrease the cell's growth rate and in turn cause it to be outcompeted in a growing population. Any subpopulation containing 'more active' mutations will therefore fail to increase in size to a significant number and thus not have a significant effect on the population's dynamics. To show this, we plot example dynamics for the toggle switch and the repressilator. In each case, we use the same parameters as for our $s_3$ simulations, except now denoting the 'intermediate' parameters with the subscript 'M*' and setting $\alpha_{\mathrm{E}} < \alpha_{\mathrm{M}^*}$.

To show the dynamics for the synthetic constructs, we plot the average concentration of each synthetic protein per cell, and the growth rate relative to the M-state of various 'states of interest': the E-state, and states with different numbers of synthetic genes having increased activity as denoted by '*'. For both constructs, the protein dynamics remain relatively stable until the the point at which the M-state starts to dominate, similar to the dynamics from

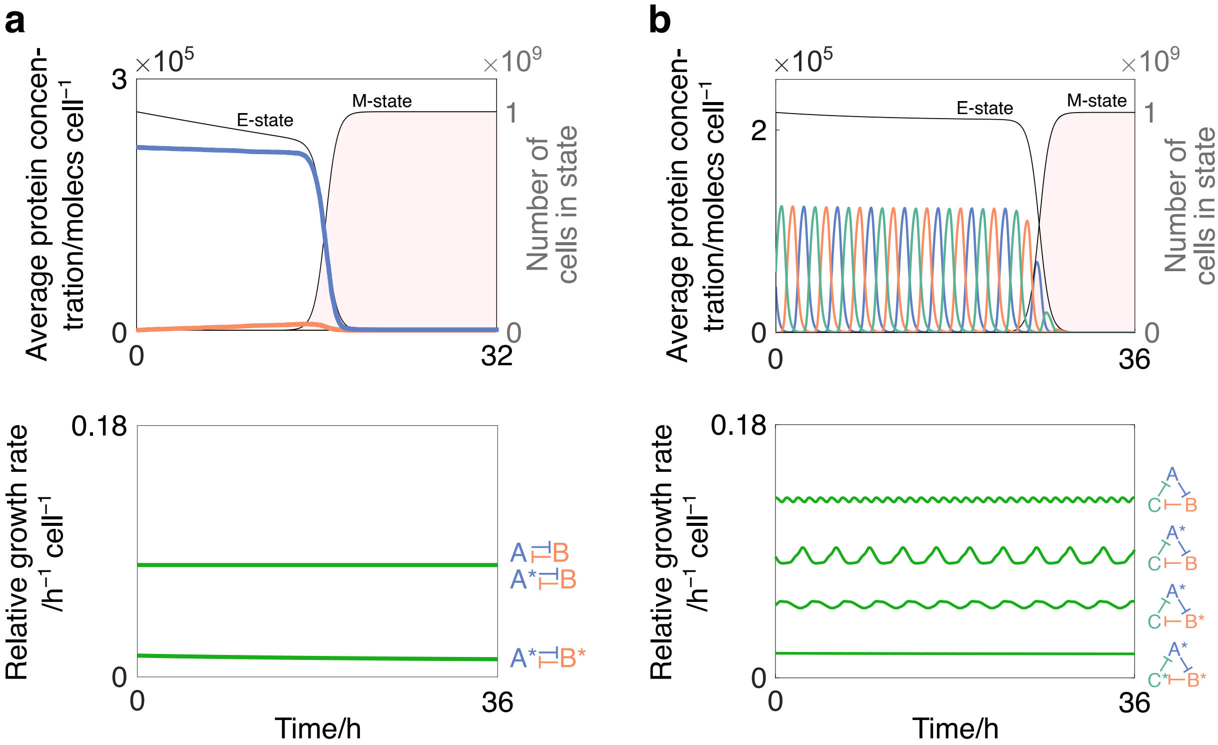


**Figure S6: Modelling synthetic constructs that can sustain function-increasing mutations.** **(a)** Toggle switch simulation. *Top*: as in Figure 3 from the main text. *Bottom*: relative growth rates are plotted for cells in three states: the E-state, the state where protein-A has increased activity, and the state where both protein-A and protein-B have increased activity. For each synthetic gene: $[\alpha_{\mathrm{E}}, z_{\mathrm{M}}, \alpha_{\mathrm{M}^*}, z_{\mathrm{M}^*}] = [10^5\,\mathrm{molecs\,h^{-1}\,cell^{-1}}, 10^{-6}, 10^6\,\mathrm{molecs\,h^{-1}\,cell^{-1}}, 5{\times}10^{-2}]$. **(b)** Repressilator simulation. *Top*: as in Figure 4 from the main text. *Bottom*: relative growth rates are plotted for cells in four states: the E-state, and states with one, two or three of the synthetic proteins having higher activity. For each synthetic gene: $[\alpha_{\mathrm{E}}, z_{\mathrm{M}}, \alpha_{\mathrm{M}^*}, z_{\mathrm{M}^*}] = [10^5\,\mathrm{molecs\,h^{-1}\,cell^{-1}}, 10^{-6}, 10^6\,\mathrm{molecs\,h^{-1}\,cell^{-1}}, 1.2{\times}10^{-2}]$. In both subplots, '*' denotes a protein associated with a function-increasing mutation.

Figure 3b and Figure 4b in the main text when only severe mutations are modelled. This can be again explained by the growth rates of the states (lower panels) where there is an increase in protein activity: increasing the activity of any combination of the synthetic genes produces no gain in growth rate, and as such no selective advantage results.

Aside from considering how mutations change the activity of a synthetic part, we note that the effect of mutations may be linked to the function of the gene itself. For purposes of generality, we have assumed that synthetic genes have no functional effect on the cell other than their drain on shared cellular resources. In other words, we have considered the effect on 'fitness' but not 'utility'. If genes were instead modelled to additionally affect core cellular processes, then the consequences of synthetic gene expression would be less clear-cut. For example, if a gene increased the rate of energy metabolism, then increasing its activity would not only drain shared cellular resources, but also increase the activity of the cell. This could easily be implemented in our model by adding a positive autoregulation term that links the concentration of synthetic protein, $H$, to the rate of energy metabolism from Equation (S2), $\epsilon(Z)$, such that the modified rate of energy metabolism, $\epsilon^*(Z)$, would be of the form:

$$\epsilon^*(Z) = \epsilon(Z) \cdot \frac{\left(\frac{H}{K_H}\right)^{h_H}}{1 + \left(\frac{H}{K_H}\right)^{h_H}}, \tag{S26}$$

where $K_H$ and $h_H$ denote the half-saturation constant and the hill coefficient, respectively. Similar effects could be applied to any of the cellular processes that our model considers, either as negative or positive regulatory effects. In turn, both fitness and utility could be modelled for any of the gene constructs we consider.

### S5 Supplementary analysis of model complexity

The complexity of our modelling framework increases significantly when more dimensions and intermediate states are added. Specifically, the total number of states and total number of transitions that need to be modelled rises sharply. While unrelated to the model applications discussed so far, analysing the complexity of frameworks has interesting links to the combinatorics branch of mathematics, and so understanding this in more detail could aid future applications and refinements of our framework. In this section, we will therefore shed some light on these themes.

#### S5.1 Total number of states

The total number of states in a system ($n$) can be calculated by raising the number states per dimension ($s$) to the power of the number of dimensions ($d$):

$$n = s^d. \tag{S27}$$

The result of this can be immediately seen from the frameworks displayed in Figure 1b in the main text. For example, the $[s_2, d_3]$ framework has $2^3 = 8$ states, while the $[s_2, d_3]$ framework has $3^3 = 27$ states. A more complex system with five mutable parts, each with four states per dimension ($[s_4, d_5]$), would then have $4^5 = 1024$ states in total.

#### S5.2 Group structures

Interesting patterns emerge when calculating the total number of transitions ($T$) within a framework for given values of $s$ and $d$. With simple frameworks, $T$ can be easily inferred, for example in the $[s_2, d_2]$ framework it is clear that the total number of transitions is four. For more complex frameworks, such as the $[s_4, d_5]$ example, a more systematic method is required. One way to approach this is to examine the geometric symmetries within the framework designs.

In each framework, cell states can be grouped according to how many outgoing transitions they have. For example, analysing the $[s_2, d_2]$ framework in Figure S7a, three groups can be made: (i) the E-state with two transitions, (ii) the two single-mutant states with one transition, and (iii) the full-mutant state with zero transitions. If the size of each group is the number of states it contains, and if the sizes are ordered according to the number of transitions that each state within that group has, then the sequence of group sizes for the $[s_2, d_2]$ framework can be represented as $\{1, 2, 1\}$. Using this logic, $T$ can be calculated as $(\mathbf{1} \times 2) + (\mathbf{2} \times 1) + (\mathbf{1} \times 0) = 4$, where the group sizes (bold numbers) are multiplied by the number of outgoing transitions that each state within that group contains. While the particular sequence for $[s_2, d_2]$ is simple, the general principle is more useful for understanding $T$ at higher values of $s$ and $d$. To formalise this idea, we define this sequence of numbers as a framework's 'group structure':

**Definition S5.1** (Group structure)**.** The sequence of numbers that represents the number of states within each 'group' for a given framework, where a 'group' contains all states with an equivalent number of outgoing transitions. This is captured in descending order according to the number of transitions that each state within a group has.

The group structures for four frameworks are visually highlighted in Figure S7. For example with the $[s_2, d_2]$ framework, the E-cell state has three outgoing transitions and each subsequent group has one fewer. The resulting group structure is therefore $\{1, 3, 3, 1\}$, and $T$ can be represented as $(\mathbf{1} \times 3) + (\mathbf{3} \times 2) + (\mathbf{3} \times 1) + (\mathbf{1} \times 0) = 12$. For each new dimension added, it emerges that the group structure is the next row of Pascal's Triangle (Figure S7a), which lists the binomial coefficients for a given number of elements (or in this case, 'dimensions'). This is equivalent to asking how many ways there are of partitioning $d$ elements into two unordered subsets of sizes $i$ and $j$.[1]

[1] As an example with three elements 'A', 'B' and 'C', there are $\binom{3}{i,j} = \frac{3!}{i!j!}$ possible pairs of sets, where $j = 3 - i$. It follows that $i = 0$ yields one pair ('nothing and ABC'); $i = 1$ yields three pairs ('A and BC', 'B and AC', 'C and AB'); $i = 2$ yields three pairs ('AB and C', 'AC and B', 'BC and A'); $i = 3$ yields one pair ('ABC and nothing').

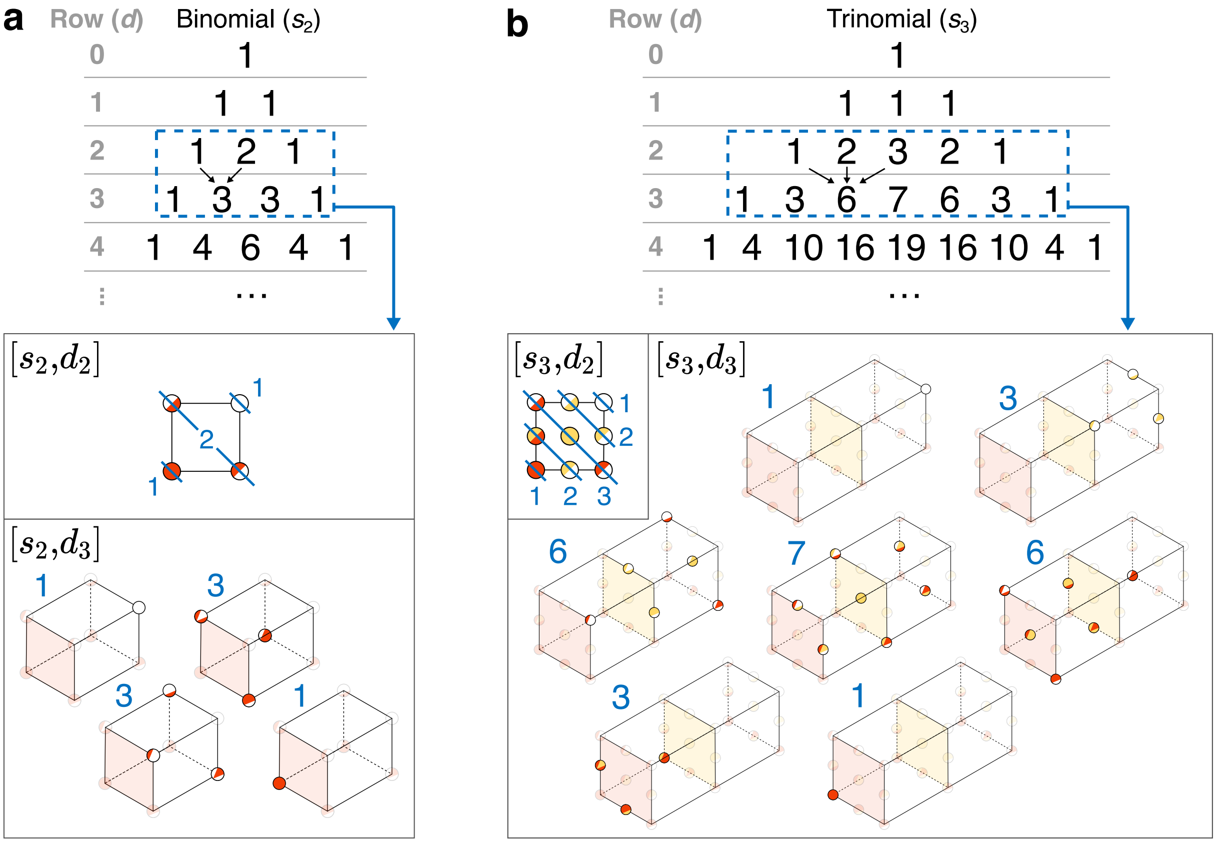


c

| | $d_1$ | $d_2$ | $d_3$ | $d_4$ | |
|---|---|---|---|---|---|
| $s_1$ | {1}— group structure<br>1 — no. transitions | {1}<br>1 | {1}<br>1 | {1}<br>1 | Monomial |
| $s_2$ | {1,1}<br>1 | {1, 2, 1}<br>4 | {1, 3, 3, 1}<br>12 | {1, 4, 6, 4, 1}<br>32 | Binomial |
| $s_3$ | {1,1,1}<br>1 | {1, 2, 3, 2, 1}<br>18 | {1, 3, 6, 7, 6, 3, 1}<br>81 | {1, 4, 10, 16, 19, 16, 10, 4, 1}<br>324 | Trinomial |
| $s_4$ | {1,1,1,1}<br>1 | {1, 2, 3, 4, 3, 2, 1}<br>48 | {1, 3, 6, 10, 12, 12, 10, 6, 3, 1}<br>288 | {1, 4, 10, 20, 31, 40, 44, 40, 31, 20, 10, 4, 1}<br>1536 | Quadrinomial |
| | 1st row of... | 2nd row of... | 3rd row of... | 4th row of... | |

Calculating number of transitions

$$81 = \text{sum}\left\{ \begin{matrix} 1 & 3 & 6 & 7 & 6 & 3 & 1 \\ \times & \times & \times & \times & \times & \times & \times \\ 6 & 5 & 4 & 3 & 2 & 1 & 0 \end{matrix} \right\} = \binom{3}{2} \cdot 3 \cdot 3^{3-1}$$

**Figure S7: Combinatorial patterns that arise by grouping state transitions.** States in any framework can be grouped according to their 'group structures' (see the text of Section S5.2). **(a)** The group structures for $s_2$ frameworks form the binomial coefficients, with frameworks of increasing dimensions forming successive rows of coefficients. Example group structures are shown for $d_2$ and $d_3$. **(b)** as in (a), but for $s_3$ frameworks and trinomial coefficients. **(c)** A table of groups structures (blue sets) and the total number of transitions (black number) for different values of $s$ and $d$. The relationship to the multinomial coefficients is shown via the grey text below each column and adjacent to each row. The total number of transitions can either be calculated systematically from the group structures or by using a general formula, as shown by the purple box.

When a third state is added to each dimension, the resulting patterns become more complex (Figure S7b). For the $[s_2, d_3]$ and $[s_3, d_3]$ frameworks, for example, the group structures are $\{1, 2, 3, 2, 1\}$ and $\{1, 3, 6, 7, 6, 3, 1\}$ which yield $T = 18$ and $T = 81$ respectively. For $s_3$, the group structure formed when adding new dimensions follows successive rows of the trinomial triangle, whose $d^{\text{th}}$ row lists the trinomial coefficients. Analogous to the binomial coefficients, this is equivalent to asking how many ways there are of partitioning $d$ elements into three unordered subsets of sizes $i$, $j$ and $k$. Extending this pattern to any value of $s$, the group structures follow rows of the 'multinomial triangle' whose $d^{\text{th}}$ row lists the multinomial coefficients of order $s$ [17].

Algebraically, a framework's group structure can be generated for any value of $s$ and $d$ by calculating the coefficients of $a$ in the polynomial expansion: $(a^0 + a^1 + \cdots + a^{s-1})^d$. For example, the expansion for the group structure of framework $[s_3, d_3]$ is $\mathbf{1}a^6 + \mathbf{3}a^5 + \mathbf{6}a^4 + \mathbf{7}a^3 + \mathbf{6}a^2 + \mathbf{3}a + \mathbf{1}$, yielding $\{1, 3, 6, 7, 6, 3, 1\}$.

##### S5.3 Total number of state transitions

While symmetries within group structures can be used to calculate $T$ for given values of $s$ and $d$, it would be more convenient to obtain a general formula for $T$. Starting with $d_1$, the number of transitions for any value of $s$ is equivalent to the number of ways of choosing any two states, as each pair of states always has a unique transition. This is equivalent to $\binom{s}{2} = \frac{s!}{2!(s-2)!} = \frac{s(s-1)}{2}$.

To calculate $T$ for $d > 1$ dimensions, it is clear that the result for one dimension needs to be increased by at least a factor $d$, however this only accounts for transitions exiting the E-cell state. To account for the other transitions, an additional factor is required which, through empirical testing, we found is likely to be $s^{d-1}$. This results in a postulated equation for $T$:

$$T = \binom{s}{2} \cdot d \cdot s^{d-1} = \frac{s^d \cdot (s-1) \cdot d}{2}. \tag{S28}$$

For example, for the $[s_3, d_3]$ framework, $T = \frac{3^3 \cdot (3-1) \cdot 3}{2} = 81$, and for the hypothetical simulation with five mutable parts and four states per dimension ($[s_4, d_5]$), $T = \frac{4^5 \cdot (4-1) \cdot 5}{2} = 7680$. The group structures and values of $T$ for the first four states and dimensions are summarised in Figure S7c.
